# Crowdsourced study of children with autism and their typically developing siblings identifies differences in taxonomic and predicted function for stool-associated microbes using exact sequence variant analysis

**DOI:** 10.1101/319236

**Authors:** Maude M David, Christine Tataru, Jena Daniels, Jessey Schwartz, Jessica Keating, Jarrad Hampton-Marcell, Neil Gottel, Jack A. Gilbert, Dennis P. Wall

**Author notes:** Corresponding Author: Dennis P. Wall, Ph.D., Department of Pediatrics, Division of Systems Medicine, Stanford University, 1265 Welch Rd, Suite X141, Stanford, CA 94305, E.

## Abstract

**Background:** The existence of a link between the gut microbiome and autism spectrum disorder (ASD) is well established in mice, but in human populations efforts to identify microbial biomarkers have been limited due to problems stratifying participants within the broad phenotype of ASD and a lack of appropriately matched controls. To overcome these limitations and investigate the relationship between ASD and the gut microbiome, we ran a crowdsourced study of families 2-7 year old sibling pairs, where one child of the pair had a diagnosis of ASD and the other child did not.

**Methods:** Parents of age-matched sibling pairs electronically consented and completed study procedures via a secure web portal (microbiome.stanford.edu). Parents collected stool samples from each child, responded to behavioral questionnaires about the ASD child’s typical behavior, and whenever possible provided a home video of their ASD child’s natural social behavior. We performed DNA extraction and 16S rRNA amplicon sequencing on 117 stool samples (60 ASD and 57 NT) that met all study design eligibility criteria,. Using DADA2, Exact Sequence Variants (ESVs) were identified as taxonomic units, and three statistical tests were performed on ESV abundance counts: (1) permutation test to determine differences between sibling pairs, (2) differential abundance test using a zero-inflated gaussian mixture model to account for the sparse abundance matrix, and (3) differential abundance test after modeling under a negative binomial distribution. The potential functional gene abundance for each sample was also inferred from the 16S rRNA data, providing KEGG Ortholog (KO), which were analyzed for differential abundance.

**Results:** In total, 21 ESVs had significantly differentially proportions in stool of children with ASD and their neurotypical siblings. Of these 21 ESVs, 11 were enriched in neurotypical children and ten were enriched in children with ASD. ESVs enriched in the ASD cohort were predominantly associated with Ruminococcaceae and Bacteroidaceae; while those enriched in controls were more diverse including taxa associated with *Bifidobacterium*, *Porphyromonas*, *Slackia*, *Desulfovibrio*, *Acinetobacter johnsonii*, and Lachnospiraceae. Exact Variant Analysis suggested that Lachnospiraceae was specific to the control cohort, while Ruminococcaceae, Tissierellaceae and Bacteroidaceae were significantly enriched in children with ASD. Metabolic gene predictions determined that while both cohorts harbor the butyrogenic pathway, the ASD cohort was more likely to use the 4-aminobutanoate (4Ab) pathway, while the control cohort was more likely to use the pyruvate pathway. The 4Ab pathway releases harmful by-products like ammonia and can shunt glutamate, affecting its availability as an excitatory neurotransmitter. Finally, we observed differences in the carbohydrate uptake capabilities of various ESVs identified between the two cohorts.

## INTRODUCTION

Autism spectrum disorder (ASD) is a heterogeneous developmental disorder affecting social and behavioral functioning in 1 out of 59 children in the United States (*1*). Recent studies have identified several environmental factors associated with ASD etiology and susceptibility, including prenatal infection (*2*), zinc deficiency (*3*), maternal diabetes (*4*), toxins and pesticides (*5*), and advanced parental age (*6*). Individuals with ASD have demonstrated a high prevalence of gastrointestinal (GI) and immunologic abnormalities pertaining to GI motility and intestinal permeability (*7*, *8*). Additionally, ASD-typified behavioral traits are more severe in children with both ASD and GI disturbances (*9*). These factors have also been shown to influence or be influenced by the intestinal microbiome (*10*), which could suggest a role for the intestinal microbiota in mediating ASD and ASD-typified behavioral traits.

The legitimacy of the proposed microbiome-ASD connection is supported by recent research on ASD phenotype mouse models and the microbiota compositions of human individuals with ASD (*11*-*18*). Hsiao *et al.* (2013) found that administrating *Bacteroides fragilis* to ASD mouse models improved ASD-typified behavioral traits by reducing anxiety, restoring communicative behaviors, and improving sensorimotor gating (*11*). Bacterial taxa, such as members of *Lactobacillus* and a genus of *Bifidobacterium,* have demonstrated microbially-induced behavioral modulation in both rats and humans (*12*-*16*). Moreover, several studies have identified microbial trends amongst the ASD population such as an increased abundance of *Clostridium* (*17*, *18*). More recently, a study involving Fecal Microbiome Transplant between neurotypical controls and children with ASD demonstrated a significant improvement in both GI and neurobehavioral symptoms following the treatment (*19*). This study particularly demonstrates a potential causative relationship between the gut microbiome and ASD symptoms. While the data on the microbiome-ASD symptom link is compelling, studies attempting to identify the specific microbes responsible have maintained small sample sizes, single time point sampling, limited phenotype scoring and sampling, and a lack of bacterial phylogenetic resolution, factors that may impact the reproducibility of the results.

The present study aims to determine the specific intestinal microbiota that associate with behavioral traits in children with ASD. We recruited families with age-matched neurotypical and ASD siblings via crowdsourcing to reach a sufficient sample size. (*20*-*26*). Recruited families had a child clinically diagnosed with ASD and a neurotypical sibling who were both between the ages of 2-7 and no more than 2 years apart in age. The crowdsourcing recruitment methodology enabled us to recruit a large cohort of families disbursed across the United States. Each family completed behavioral and dietary questionnaires online and collected a stool sample from each child at home via sampling kits shipped to each family by the research team. This approach facilitated the collection of diverse and pertinent metadata regarding allergies, diet, supplement usage, gastrointestinal abnormalities, gestational age, and antibiotic and probiotic treatment (*22*-*25*, *27*). We confirmed the self-reported autism diagnosis of each child by leveraging validated machine-learning classification tools that assess ASD-typified features obtained from parent reports and home video showcasing social interactions (*28*-*33*).

## RESULTS

### Crowd Sourcing Recruitment and Participant Demographics

Between March 2015 and September 2017, 20,478 unique users visited our study website, 1,953 were electronically screened for eligibility by survey, and 297 of them met our study inclusion criteria. 194 users electronically consented to participate, and 164 began responding to the online surveys. Of 164 participants, 100 completed the online component and were mailed sampling kits. 71 families, or parents of 142 sibling pairs, completed the online and at-home sampling procedures for the study, and 117 child-subjects (60 ASD and 57 NT) met all eligibility criteria, including the required confirmation of diagnosis obtained from the MARA and video classifier, when submitted. Of the 117 child-subjects, there were 55 sibling pairs, two sibling pairs were accompanied by a third sibling with autism, and 5 were singleton samples.

The ASD cohort comprised 72% male participants (n = 43), as compared to 55% of the NT cohort (n = 27). Dietary and lifestyle questionnaires were completed, in entirety, for 106 of the 117 participants. Among the 106 child-subjects, 66% (n = 79) identified as Caucasian, 7.5% (n = 8) identified as Asian or Pacific Islander, 3.8% (n = 4) identified as African American, and 7.5% (n = 8) identified as Hispanic (participants were also given the option to select more than one identifying ethnicity, not reported here). Participant age was not significantly different between the ASD and NT cohorts. Additional demographic data are in Supplementary Information SI 1.

### ASD Diagnosis Confirmation using the Mobile Autism Risk Assessment (MARA) and Video Classifier

All child-subjects with ASD that completed the MARA and of these 29 provided scorable video (including one family with two siblings with ASD and one NT sibling meeting the age criteria). There was a 100% agreement in class assignment between the MARA and the video classifier in all 37 cases (SI 2). In 12 instances, the output from either or both classifier (2 supported by the video classifier) did not confirm the parent-reported ASD diagnosis. These participants were therefore excluded from analysis. The results reported hereafter include only the remaining 60 child-subjects with confirmed ASD.

### Diet Differences between Children with ASD and Neurotypical Siblings

We found three categorical factors (supplements, dairy intolerance, and dietary restrictions) to be significantly different between the two cohorts according to a chi-square test (Table 1). Nutritional/herbal supplements showed significant differences between the two cohorts, with 63.6% (n = 35) ASD child-subjects taking an herbal supplement as compared to 35.3% (n = 18) NT child-subjects (qval 2.3e-2). Dairy intolerance was also more prevalent in the ASD cohort (n = 1 NT child-subjects versus n = 16 ASD child-subjects) (qval 2.6e-3), which correlates with a statistically significant deviation in the frequency of consumption of both milk/cheese and milk substitutes: only 33.3% (n = 18) of ASD child-subjects consumed milk/cheese on a “regular” or “daily” basis as compared to 72 % (n = 36) of NT child-subjects (qval 2.3e-3). Finally, gluten intolerance was found to be more prevalent in the ASD cohort (qval 3.5e-4). Additionally, n = 20 ASD child-subjects had other special dietary restrictions apart from dairy and gluten constraints, compared to only n = 6 NT child-subjects (qval 2.3e-3). Refer to Table 1 and Supplementary Information SI 1 for a summary of the remaining reported data.

**Table 1:** Clinical Characteristics for ASD and NT Participants with significant difference between the cohorts.

|  | ASD(n=60) | % | NT(n=57) | % | Adjusted p-value |
| --- | --- | --- | --- | --- | --- |
| Gender<sup>1</sup> |  |  |  |  |  |
| Male | 43/60 | 71.7% | 27/49 | 55.1% | 2.3e<sup>-2</sup> |
| Female | 17/60 | 28.3% | 22/49 | 44.9% |  |
| Gluten Intolerance<sup>2</sup> |  |  |  |  |  |
| Yes | 22/59 | 34.4% | 1/56 | 1.8% | 3.5e<sup>-4</sup> |
| No | 42/59 | 71.2% | 55/56 | 98.2% |  |
| Nutritional/herbal supplement<sup>3</sup> |  |  |  |  |  |
| Yes | 35/55 | 63.6% | 18/51 | 35.3% | 2.3e<sup>-2</sup> |
| No | 20/55 | 36.4% | 33/51 | 64.7% |  |
| Special Diet Restrictions<sup>4</sup> |  |  |  |  |  |
| Yes | 20/54 | 37.0% | 6/51 | 11.8% | 2.3e<sup>-2</sup> |
| No | 34/54 | 63.0% | 45/51 | 88.2% |  |
| Dairy Intolerance |  |  |  |  |  |
| Yes | 16/60 | 26.7% | 1/57 | 1.8% | 2.6<sup>-6</sup> |
| No | 44/60 | 73.3% | 56/57 | 98.2% |  |
| Consumption of at least 2 servings of milk or cheese a day<sup>6</sup> |  |  |  |  |  |
| Never | 23/54 | 42.6% | 6/50 | 12.0% | 2.3<sup>-3</sup> |
| Rarely(a few times a month) | 8/54 | 14.8% | 2/50 | 4.0% |  |
| Occasionally(1-2/week) | 5/54 | 9.3% | 6/50 | 12.0% |  |
| Regularly(3-5/week) | 7/54 | 12.9% | 18/50 | 36.0% |  |
| Daily | 11/54 | 20.4% | 18/50 | 36.0% |  |

<sup>1</sup> missing 8 NT gender responses
<sup>2</sup> missing 1 NT and 1 ASD responses
<sup>3</sup> missing 6 NT and 5 ASD
<sup>4</sup> missing 6 NT and 6 ASD
<sup>6</sup> missing 7 NT and 6 ASD

### Dietary and Lifestyle Habits Influencing the Microbial Community

SI 3 detailed the five variables that seems to significantly influence the microbial community: Probiotics, Multi-vitamin, sugary sweet, olive oil and sequecing batch. Constraint PcoA were used to identify the ESVs for which the abundance was the most influenced by these variables (SI 4).

### Similarity between Sibling Lifestyles

As hypothesized, sibling lifestyles, as measured by dietary choices, supplement intake, exercise, allergies, and other factors, were significantly more similar to each other than to other participants (p << .01). We used the Euclidean distance between lifestyle description vectors to perform a permutation test (999 permutations), which confirmed the similar sibling lifestyle hypothesis. (SI 5).

### Reported Gastrointestinal Symptoms

As reported above, we observed significant differences in gluten and dairy intolerances, which imply greater propensity for GI abnormalities among the ASD cohort. We did not, however, observe any significant differences between our cohorts regarding the reported gastrointestinal motility (SI 6) nor the frequency distribution of bowel movements (two-sample Wilcoxon signed-rank, p-value = 0.8313). Grouping samples into stool categories “Frequent”(> once a day), “Typical” (once a day) and “Sparse”(< once a day) did not result in any significant interdependence of phenotype and bowel movement frequency. When samples were agglomerated into two categories, typical bowel movement (one per day) and abnormal bowel movement (less or more than one per day), we did observe a slight, though not statistically significant, trend in the ASD cohort towards increase abnormal bowel movement frequency (chi-square p = 0.17).

### Microbial Alpha-Diversity

We calculated the phylogenetic diversity (PD) and Shannon diversity metric for each sample (*34*). Grouping diversity measurements into “low”, “medium” and “high” categories based on observed standard deviation, we see a significant relationship between phenotype and diversity (fisher-exact p = 0.01), with high diversity associated with ASD (Figure 1). However, performing a rank sum test using each metric, we found no significant difference between cohorts. Although the variance of diversity (distribution of scores) in the ASD cohort was significantly greater than the NT cohort (bootstrap p <.001; Figure 1). Shannon diversity was also significantly related to bowel movement quality (fisher-exact p = .02), with low diversity associated with diarrhea, but not significantly related to bowel movement frequency (fisher-exact p = .17)

**Figure 1:**
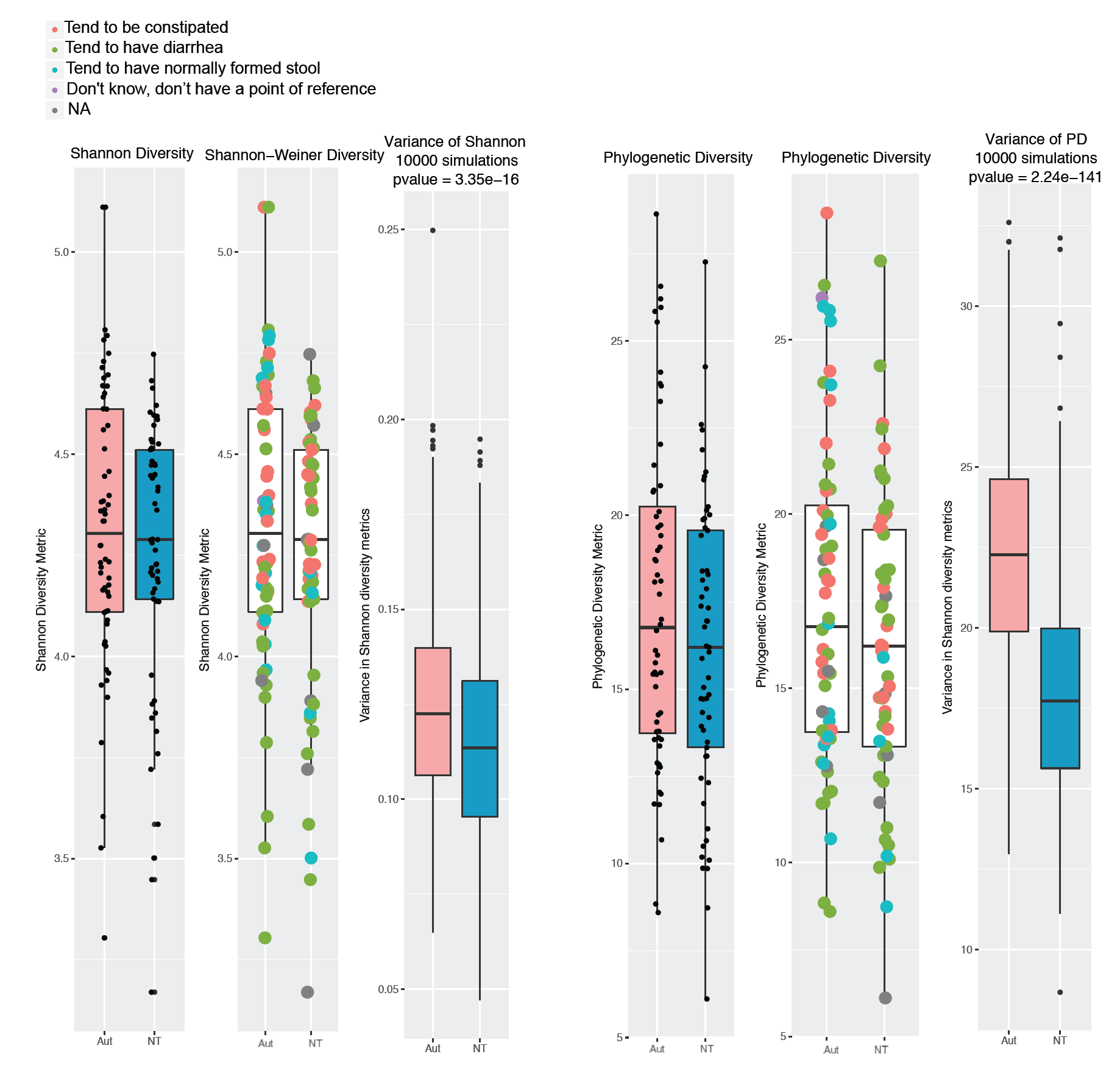
Phylogenetic Diversity and Shannon Diversity used as estimators of microbial alpha-diversity. The variance of diversity (distribution of scores) in the ASD cohort was significantly greater than the NT cohort (bootstrap p < .001) with both Diversity Estimator. Shannon diversity was also significantly related to bowel movement quality (fisher-exact p = .02), with low diversity associated with diarrhea, but not significantly related to bowel movement frequency (fisher-exact p = .17).

### Permutation Test on Sibling Pairs to determine ESVs that differentiate between ASD and NT

Ten ESVs were determined to be differentially abundant between sibling pairs (ASD vs. NT) as determined by a permutation test with FDR correction (Table 2; SI 7). The mean differential abundance drawn from the null distribution was never more extreme than the actual differential abundance mean, and all p-values were 0, and increased to 9.36 × 10^−3^ upon correction (see distribution plot in SI H). The genera *Aggregatibacter*, *Anaerococcus* and *Oscillospira* were significantly enriched in the ASD cohort, while *Porphyromonas*, *Slackia*, *Desulfovibrio*, *Clostridium colinum*, and *Acinetobacter johnsonii* were enriched in the NT cohort.

**Table 2:** Candidate 16S biomarkers enriched and depleted in the autism cohort and their annotation. This table indicates the Exact Sequence Variants (ESVs) identified using 3 analysis methods: Permutation Test on Sibling Pair Differentials (Pair), Differential Ribosomal Analysis Based on the Negative Binomial Distribution (Deseq2), and Zero Inflated Gaussian Analysis (ZIG). The annotation was performed using Ribosomal Database Project’s naive Bayesian classifier with GreenGenes database 13.8. A blast was also performed using IMG’s most recent database (January 2018), and the perfect match (100% similarity on full length of the query) is indicated in the last column.

|  | ESV | Phylum | Class | Order | Family | Genus | Species | Analysis | pvals_adj | IMG genome perfect alignment (number of hits) |
| --- | --- | --- | --- | --- | --- | --- | --- | --- | --- | --- |
| ASD CANDIDATE BIOMARKERS | ESV1 | Bacteroidetes | Bacteroidia | Bacteroidales | *Bacteroidaceae* | *Bacteroides* |  | DEseq2 | 2.34E-14 | *Bacteroides vulgatus* |
|  | ESV2 | Firmicutes | Clostridia | Clostridiales |  |  |  | Pair | 8.92E-03 | - |
|  | ESV3 | Firmicutes | Clostridia | Clostridiales | *Ruminococcaceae* |  |  | Pair | 8.92E-03 | - |
|  | ESV4 | Firmicutes | Clostridia | Clostridiales | *Ruminococcaceae* | *Oscillospira* |  | Pair | 8.92E-03 | - |
|  | ESV5 | Firmicutes | Clostridia | Clostridiales | *Tissierellaceae* | *Anaerococcus* |  | Pair | 8.92E-03 | *Anaerococcus senegalensis* |
|  | ESV6 | Proteobacteria | Gammaproteobacteria | Pasteurellales | *Pasteurellaceae* | *Aggregatibacter* |  | Pair | 8.92E-03 | *Haemophilus pittmaniae*, *Aggregatibacter* |
|  | ESV7 | Firmicutes | Clostridia | Clostridiales | *Ruminococcaceae* | *Ruminococcus* |  | ZIG | 1.09E-05 | - |
|  | ESV8 | Bacteroidetes | Bacteroidia | Bacteroidales | *Bacteroidaceae* | *Bacteroides* | *uniformis* | ZIG | 4.88E-04 | *Bacteroides uniformis* (6), *Bacteroides* sp. 4_1_36 (1) |
|  | ESV9 | Firmicutes | Erysipelotrichi | *Erysipelotrichales* | *Erysipelotrichaceae* | *Holdemania* |  | ZIG | 2.20E-02 | *Holdemania filiformis* |
|  | ESV10 | Firmicutes | Clostridia | Clostridiales | *Clostridiaceae* | *Clostridium* | *celatum* | ZIG | 2.70E-02 | *Clostridium saudiense* <sup>1*</sup> |
| NEUROTYPICAL CANDIDATE BIOMARKERS | ESV11 | Firmicutes | Clostridia | Clostridiales | *Lachnospiraceae* | *[Clostridium]* | *colinum* | Pair | 8.92E-03 | - |
|  | ESV12 | Bacteroidetes | Bacteroidia | Bacteroidales | *Porphyromonadaceae* | *Porphyromonas* |  | Pair | 8.92E-03 | - |
|  | ESV13 | Actinobacteria | Coriobacteriia | *Coriobacteriales* | *Coriobacteriaceae* | *Slackia* |  | Pair | 8.92E-03 | *Slackia piriformis* |
|  | ESV14 | Proteobacteria | Deltaproteobacteria | *Desulfovibrionales* | *Desulfovibrionaceae* | *Desulfovibrio* |  | Pair | 8.92E-03 | *Desulfovibrio fairfieldensis* (3), *Desulfovibrio* sp. 3_1_syn3 (3), *Desulfovibrio* sp. 6_1_46AFAA (1) |
|  | ESV15 | Proteobacteria | Gammaproteobacteria | *Pseudomonadales* | *Moraxellaceae* | *Acinetobacter* | *johnsonii* | Pair | 8.92E-03 | *Acinetobacter johnsonii* (18), *Acinetobacter johnsonii* (4), *Acinetobacter schindleri* |
|  | ESV16 | Firmicutes | Clostridia | Clostridiales | *Lachnospiraceae* | *Lachnospira* |  | ZIG | 2.12E-03 | *Eubacterium eligens* <sup>2*</sup> |
|  | ESV17 | Firmicutes | Clostridia | Clostridiales | *Lachnospiraceae* |  |  | ZIG | 9.73E-03 | - |
|  | ESV18 | Firmicutes | Clostridia | Clostridiales | *Lachnospiraceae* |  |  | ZIG | 3.52E-03 | - |
|  | ESV19 | Firmicutes | Clostridia | Clostridiales | *Lachnospiraceae* | *[Ruminococcus]* |  | ZIG | 1.64E-02 | *Ruminococcaceae* bacterium GD1 (2), *Ruminococcus* sp. DSM 100440 (1), *Clostridiales* bacterium VE202-14 (1), *Sellimonas intestinalis* BR72 (1) |
|  | ESV20 | Firmicutes | Clostridia | Clostridiales | *Lachnospiraceae* | *Coprococcus* | *catus* | ZIG | 3.00E-02 | *Coprococcus catus* |
|  | ESV21 | Actinobacteria | Actinobacteria | *Bifidobacteriales* | *Bifidobacteriaceae* | *Bifidobacterium* |  | ZIG | 4.87E-02 | *Bifidobacterium pseudocatenulatum* (9), *Bifidobacterium catenulatum* (6), *Bifidobacterium gallicum* (4), *Bifidobacterium* |

ESVs with taxa specific to the ASD cohort
ESVs with taxa specific to the NT cohort
<sup>1\*</sup> *Clostridium celatum* being the second hit with 232/233 pb
<sup>2\*</sup> *Lachnospira* first hit 221/233

### Models to Maximize the Likelihood of Detecting Low Abundance Species

We implemented a mixture model using a zero-inflated Gaussian (ZIG) distribution of mean group abundance for each ESV in metagenomeSeq (*35*), in order to quantify the fold change in taxa between the ASD cohort and the NT cohort. This analysis again revealed 10 ESVs differentially present in the two cohorts: four were enriched in the ASD cohort (Table 2), and six were enriched in the NT cohort. The NT cohort was enriched in the Lachnospiraceae (five of six ESVs), including *Coprococcus catus*, *Clostridium colinum*, and the genera *Bifidobacterium.* In comparison, the ASD cohort was enriched in *Ruminococcus* and *Holdemania,* as well as the species *Bacteroides uniformis* and *Clostridium celatum.*

We also used a Negative Binomial Distribution Analysis to Identify *ESV between ASD and NT*, through which we identified a single ESV, from the genus *Bacteroides* (ESV1), enriched in the ASD cohort. Among the aforementioned statistical analyses, the microbial genus types identified in more than one statistical abundance test in the ASD cohort included the family *Ruminococcaceae* (by three ESVs including the genera *Oscillospira* and *Ruminococcus*), and the genus *Bacteroides* (by two ESVs). The NT cohort presents six ESVs belonging to the family *Lachnospiraceae*.

### Functional Profile Prediction

The software Piphillan predicted ∽6900 active KEGG Orthologs (KO) that were part of ∽170 metabolic pathways as defined by KEGG Brite. Overall, we were able to associate 105 ESVs with full genome annotations. From the predicted KOs that were present in these genomes, we observed 17 predicted metabolic pathways with significantly differential abundance between ASD and NT. Two pathways were significantly enriched in the ASD cohort: Flagellar assembly (ko02040), and Aminoacyl-tRNA biosynthesis (ko00970) (Figure 2). Fifteen pathways were significantly enriched in the NT cohort, including Butanoate metabolism (ko00650), Propanoate metabolism (ko00640), Sulfur metabolism (ko00920), Phosphotransferase system (ko02060), and microbial metabolism in diverse environments (ko01120). A full list of significantly differential pathways is in Figure 2.

**Figure 2:**
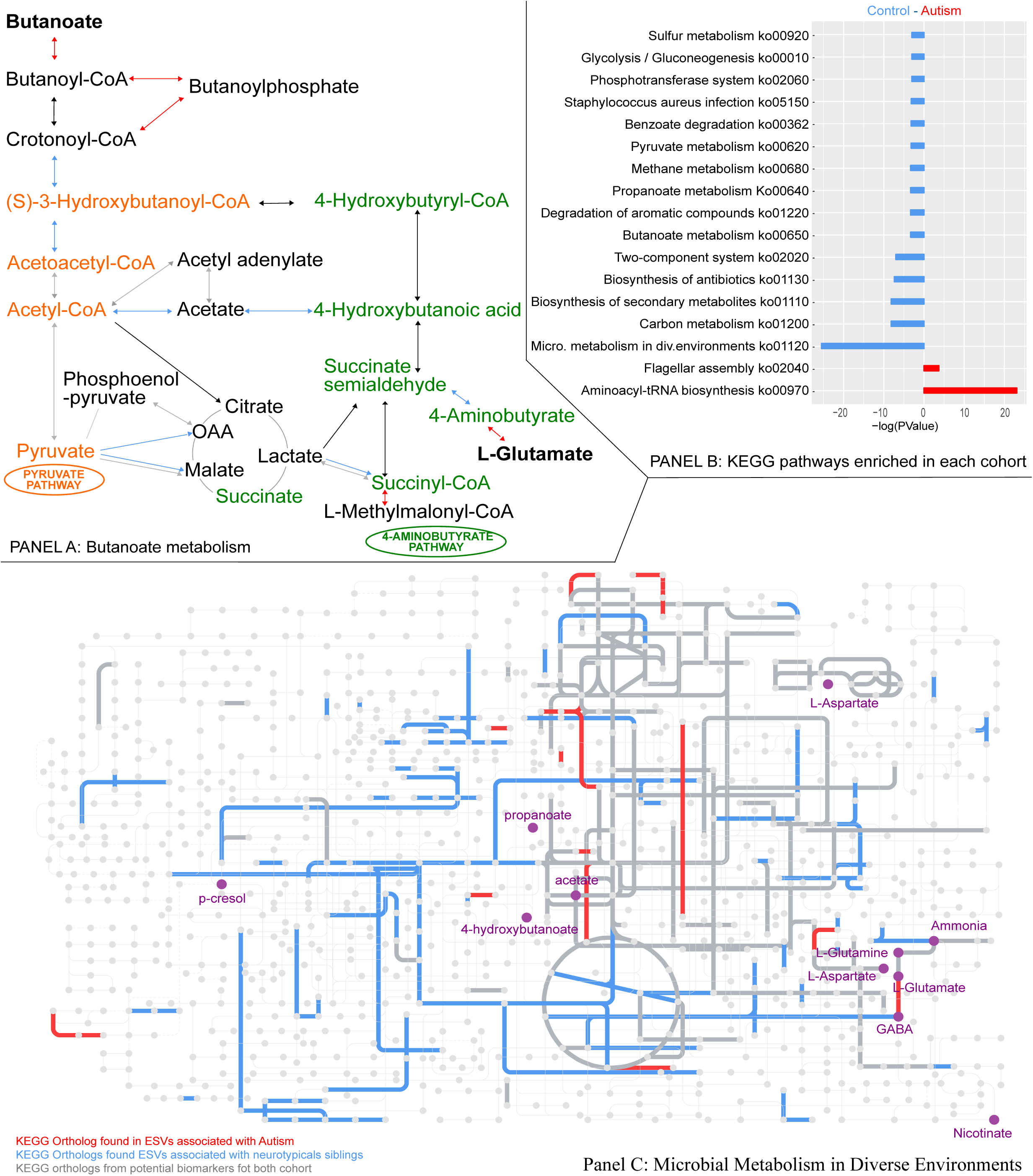
Pathway analyses derived from Exact Sequence Variants Analysis. **Panel A: Butanoate metabolism:** Detailed analysis of the butanoate pathway, the color of the arrow reflecting the cohort in which the ESV carrying the KEGG ortholog was detected **Panel B: KEGG pathways enriched in each cohort:** List of the 17 pathways enriched in the Gene Set Enrichment Analysis using genomes and abundances estimate from the ribosomal analysis **Panel C: Microbial Metabolism in Diverse Environments:** Detailed analysis of the pathway microbial metabolism in diverse environments and metabolites of interest for the gut-brain axis interaction.

## DISCUSSION

### Crowdsourcing Recruitment

By targeting the Internet-active autism community, we were able to crowdsource study subject recruitment and reach our targeted sample size for each cohort in a short amount of time. This methodology allowed us to collect data efficiently and effectively and to recruit participants from diverse geographical areas.

### Lifestyle, Dietary Practices and GI symptoms

This study highlights ASD biomarker candidates while minimizing the impact of confounding environmental factors by crowdsourcing recruitment of ASD child-subjects who have age-matched NT siblings to act as study controls. By recruiting only sibling pairs who are within 2 years of age of one another, living in similar home environments, and eating similar diets (see SI 5), we successfully controlled for diet and lifestyle among our two cohorts. Using this approach of working with sibling cohorts, other studies have also showed very similar microbial structure between the two cohorts (*36*) to the point of not being able to identify taxa specific to one or the other cohort.

We observed no overlap between factors that heavily influence the microbial structure (SI 3) and the factors that were significantly different between the ASD and the NT cohorts (Table 1). Therefore, it is unlikely that the ESVs identified as ASD or NT biomarkers were differentially abundant due to diet or lifestyle as confounding influences.

We did not observe significant differences in the GI motility or stool quality between cohorts, however, there was an increased prevalence of dairy and gluten sensitivities among ASD child-subjects which may imply a propensity for GI distress. Dairy and gluten sensitivities have been previously found to be associated with children with ASD (*37*-*39*). We also observed an increase in special dietary restrictions among our ASD child-subjects, which could imply that many parents had already implemented limitations to their child’s diet to alleviate any potential or previously identified gastrointestinal issues. Perhaps due to high parent involvement, we did not observe the expected differences in GI motility or gastrointestinal abnormalities. Failure to detect systematic GI distress in ASD could also be attributed to differences in study populations, as older siblings with wider intervals in age as were included in Son *et al*. are more likely to have more variable lifestyles and therefore greater differences in GI state.

### Microbial Community Diversity

There is much debate in the literature as to whether microbial diversity is significantly different in children with ASD versus neurotypically developing children. While we found no significant rank sum relationship between alpha diversity of the microbiota and ASD diagnosis, we observed a significant relationship when considering samples as “High”, “Medium”, or “Low” diversity. This finding implies that more or less gut microbiome diversity does not directly translate to more or less benefit in the case of ASD, but rather that diversity should be viewed more broadly as a general contributing metric. The variance in diversity scores was significantly greater in the ASD cohort compared with the NT cohort, which may explain some of the discrepancies seen in smaller cohort studies such as those conducted by Finegold et al. (2010), Kang et al. (2017), and Hsiao et. al. (2013), which respectively report increased, decreased, and unchanged microbial diversity in an ASD cohort. Notably, in our ASD cohort, low diversity seemed strongly related to parent-reported diarrhea occurrences. Therefore, it is possible that studies that specifically enrich for significant gut abnormalities when recruiting ASD subjects may unwittingly enrich for decreased ASD microbial diversity. This finding of greater variance in the diversity of ASD microbiota suggests that perhaps the ‘Anna Karenina principle’ is at work in ASD, whereby there are more ways to be dysbiotic than non-dysbiotic, hence it is more probable to identify a greater range of alpha diversity scores in dysbiotic individuals (*40*).

### Differential abundance analysis of microbial species

This study was designed to include controls that match as exactly as possible the lifestyle and environment of the autism samples in order to allow for better reproducibility and more robust association exploration. There is a concern that fecal bacteria from ASD cases could transfer to neurotypical control siblings, thus obscuring signal and not allowing us to observe differences between the ASD typified gut and the neurotypical gut (*40*). This may be the case; however, this type of contamination only serves to obscure signal, not to create spurious results. Therefore, while there may be true biological associations not reported here, the associations observed from this cohort are not likely to be spurious.

To improve the taxonomic resolution of the analyses, this study relied exclusively on ESVs, meaning the taxonomic comparisons were performed without any clustering of 16S rRNA amplicon sequences (Table 2) (*41*). Importantly it also produces single sequence variants which can be reproducibly detected between studies and across sequencing runs, reducing potential batch effects and improving future meta-analyses (*42*).

It should be noted that while increased resolution can help improve reproducibility, no singular species, genus, or family can be considered homogeneous, and the functioning of any microbial group can vary widely based on circumstances including most recent diet, influence of other microbial community members, and strain level variation. The same species can manifest different effects on host physiology, but we report here cohort-wide associations that appear robust in a large subset of our population.

### ESVs enriched in the ASD cohort

We provided in Table 3 & Table 4 a comprehensive list of biological information and information related to other reports on each bacterial associations with ASD and other pertinent phenotypes. Our findings agree overall with literature that states that *Bacteroides* genus and families such as Erysipelotrichaceae and Clostridiaceae (member of Clostridial cluster I) have already been widely reported as enriched in ASD (see Table 3). We also observed that members of Clostridial cluster IV (genus *Ruminococcus)*, and ESV5 which belongs to the family Pasteurellaleae, are both enriched in the ASD cohort. This entire family was previously reported as being depleted in ASD participants (*43*); this discrepancy could be explained by the study’s aggregation of all Pasteurellaleae, while we implicate a single member of the family. Pasteurellaleae was also detected as one of the most abundant bacterial family in children with developmental disabilities in Japan (*44*), supporting its potential association with atypical behavioral phenotypes. Finally, we also found that the genus *Anaerococcus* (ESV5) was enriched in the ASD cohort which to our knowledge, has not yet been reported in previous research literature.

**Table 3:**
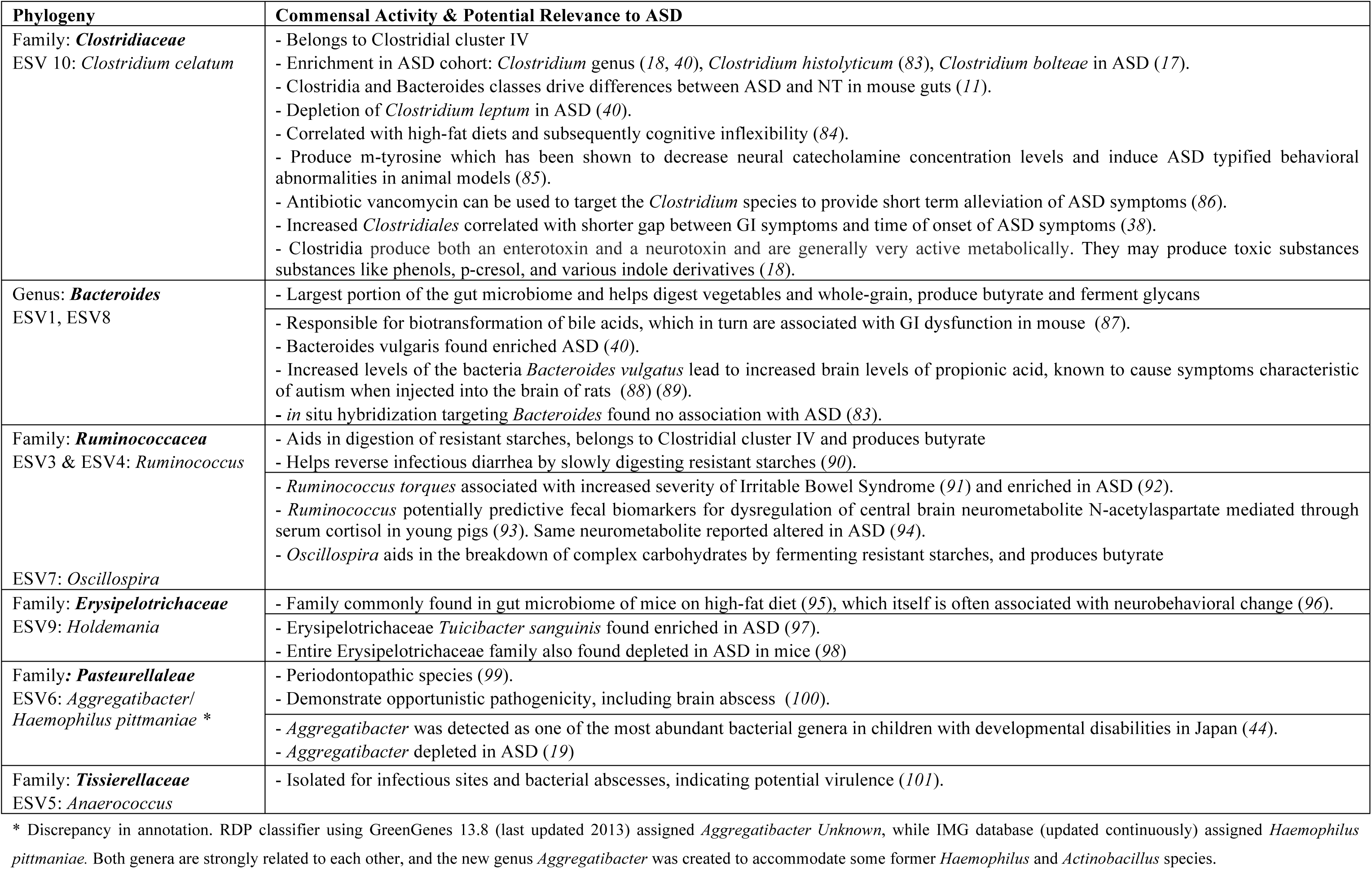
Taxa Enriched in the ASD Cohort:

**Table 4:** Taxa Enriched in the Neurotypical Cohort:

| Phylogeny | Commensal Activity & Potential Relevance to ASD |
| --- | --- |
| Family: ***Lachnospiraceae***<br>ESV17 & ESV18<br>ESV16: *Lachnospira*<br>ESV20: *Coprococcus catus*<br>ESV19: *Ruminococcus* | - Member of Clostridium cluster XIV (45).<br>- Help digest fiber and produces butyrate (102).<br>- Family shown depleted in inflammatory bowel disorder (103).<br>- Genus *Coprococcus* found depleted in ASD (13, 43)<br>- Plethora of *Lachnospiraceae* depletion in ASD suggests lack of bacterial taxa important for carbohydrate degradation (39). |
| Family: ***Desulfovibrionaceae***<br>ESV14: *Desulfovibrio* | - *Desulfovibrio* genus potentially key influential organisms in ASD (69) as children with ASD commonly have low blood levels of sulfur and high urinary excretion.<br>- *D. pigers*, *D. desulfuricans*, and *D. intestinalis* found enriched in severe ASD (40), but multiple hypothesis testing correction not performed.<br>- No association found at the genus level after multiple hypothesis testing correction (13).<br>- Increased abundance correlated with improvement of GI and improvement of behavioral ASD symptoms after microbial transfer therapy (19). |
| Family: ***Bifidobacteriaceae***<br>ESV21: *Bifidobacterium* | - Provides protection from pathogenic infections (104).<br>- assists in normalization of gut permeability and inhibits inflammatory cytokine IL-10 (16)<br>- Aid in prevention of diarrhea, reduce food allergies, and help digest lactose (105).<br>- Found depleted in ASD (40, 106)<br>- Found increased in ASD after microbiota transfer therapy that correlated with improvement of GI and behavioral symptoms (19). |
| Family: ***Porphyromonadaceae***<br>ESV12 | - Produces butyrate (107)<br>- Genus *Porphyromonas* harbors several species known to be pathogens for oral cavity, namely *Gingivalis* (108).<br>- *Gingivalis* produces a unique capsular polysaccharide that triggers Toll-like Receptor 2 dependent anti-inflammatory mechanisms in autoimmune encephalomyelitis mice (model for Multiple Sclerosis) (109, 110). |
| Family: ***Coriobacteriaceae***<br>ESV13: *Slackia* | - relatively little physiological information available about the *Slackia* genus (111). |
| Family: ***Moraxellaceae***<br>ESV15 *Acinetobacter johnsonii* | - Has been characterized in soil and water for its catabolic property in degrading aromatic compounds (112).<br>- Multiple strains of this species have also been reported to develop multi-antibiotics resistance, most likely through horizontal transfer reshuffling (114).<br>- Has capacity to proliferate on butyrate as a sole carbon source (115). |

\*We will not discuss the results related to the ESV2 as it was not possible to assign any taxonomy beyond the order level.

### ESVs depleted in the ASD cohort

Our analysis identified five ESVs belonging to the Lachnospiraceae family that are depleted in the ASD cohort and enriched in the neurotypical cohort. This family overall has already been reported as associated with autism phenotype (see Table 3). Of these ESVs, the RDP classifier was only able to assign two species names, and we were only able to identify three genomes carrying similar ribosomal sequences (Table 2), indicating that we may have identified novel variants from our analyses (*45*, *46*).These homogeneous phylogenetic groups of ESVs seem especially interesting as they cluster near each other on the 16S rRNA phylogenic tree (Figure 3). Additionally, members from the *Clostridial* cluster IV were associated with ASD, while members from the *Clostridial* cluster XIVa were associated with the control cohort. We could hypothesize that these two families exhibit metabolic pathways that are distinct among their functional redundancies. As microbes from this genus are some of human gut-associated microbiomes main butyrate producers, we examine the differences in butyrate production pathways of these two clusters in our pathway analysis below.

**Figure 3:**
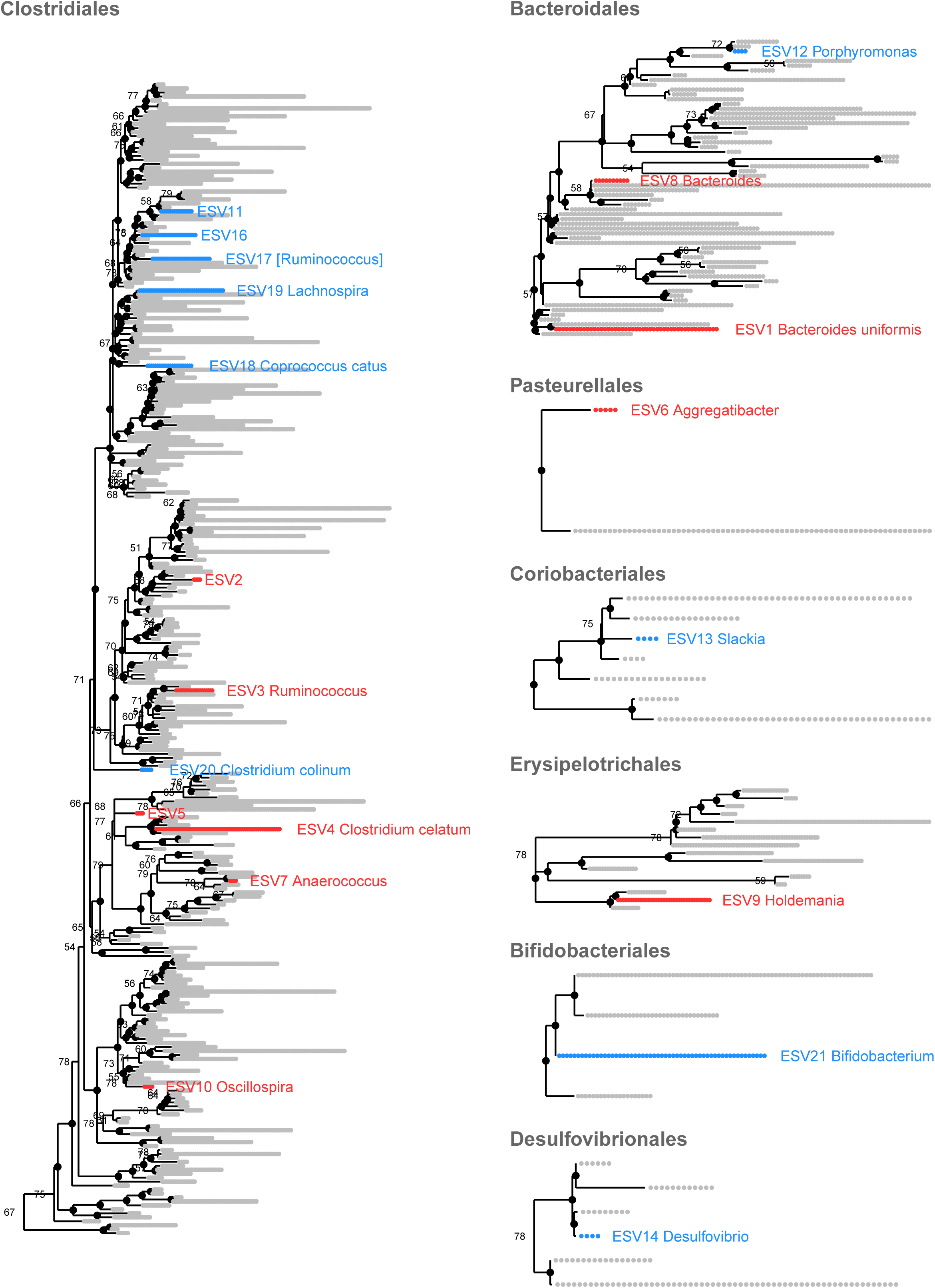
Selected order from V4 ribosomal derived phylogenetic tree. Ribosomal derived phylogenetic tree (V4 alignments) of the order carrying Exact Sequence Variants (ESV) of interest. The ESV in blue are enriched in Neurotypical cohort and the ones in blue in the Autism cohort.

The genera *Desulfovibrio* and *Bifidobacterium* were also depleted in the ASD cohort, consistent with results from at least 3 other studies (see table 3). *Bifidobacterium* has been characterized for its ability to normalize gut permeability (*16*), and lack of this genus has been hypothesized to facilitate translocation of harmful microbial metabolites from the gut to the blood. Our analysis also pinpoints the possible importance of the genus *Slackia* in the neurotypical cohort, which was depleted in our ASD cohort. Finally, we identified two more families depleted in the ASD cohort, Porphyromonadaceae and Moraxellaceae, which respectively produce butyrate or use it as the sole source of carbon.

### Pathway analysis

Using the pipeline Piphillin (*47*) to infer KOs from our ESVs, and performing a GSEA using the KOs, we identified 17 predicted pathways associated with either cohort. Predicted pathway abundance in this analysis is defined by the relative potential capacity of a present bacterial genome to produce any active enzymes in a given biological pathway. It should be noted that connecting enriched predicted pathways to the potential biomarker ESVs reported can be tenuous, as Piphillin was only able to match 6 of our 21 markers to full genomes and extract their associated KOs. Notable differentially predicted pathways include (1) butanoate metabolism, glycolysis and pyruvate metabolism, (2) propanoate metabolism, (3) sulfur metabolism, (4) aminoacyl-tRNA biosynthesis, (5) the phosphotransferase system, and (6) microbial metabolisms in diverse environments. Additionally predicted pathways comprised more general functions (e.g., biosynthesis of antibiotics, biosynthesis of secondary metabolites, carbon metabolism, and two-component system pathways) or were only detected in one ESV biomarker (e.g. flagellar assembly).

### Butyrate Production Pathway

The potential role of short chain fatty acids (SCFAs) in autism has been discussed in multiple studies. Wang et al. reported elevated SCFA concentration in children with ASD (*48*), while two other studies reported the opposite trend when looking at total SCFAs (*9*, *49*). MacFabe *et al*. found that intravenous administration of the SCFA propionate induced ASD typified behavior in mouse models, though it is likely that propionate injected intravenously may have a different effect compared with propionate originating from GI-microbial fermentation (*50*). Butyrate, in particular, has been proposed as a potential major mediator of the gut-brain axis either through modulation of the density of cholinergic enteric neurons through epigenetic mechanisms, or through direct modulation of the vagus nerve and hypothalamus (*51*). In our cohort, we observe an enrichment of microbial genomes capable of butyrate metabolism, implying that butyrate production and consumption pathways, in the stool-microbiome of NT participants (Figure 2). Butyrate production pathways in commensal microbial species and pathogens are thought to have evolved divergently; there are 4 pathways for butyrate production each branching from a different initial substrate: Pyruvate, 4-aminobutyrate, Glutarate, and Lysine (*52*). The by-products and influences of these major butyrogenic pathways could be relevant to host physiology. In our cohort, the predicted bacterial genetic potential in NT samples showed an enrichment for KOs associated with butyrate production from pyruvate, while in the ASD samples, the predicted functional potential was enriched for butyrate production via the 4-aminobutanoate (4Ab) pathway (Figure C). 4Ab is a neurotransmitter and its biosynthesis can directly interfere with the amount of available glutamate (*52*). This pathway can also potentially release harmful by-products such as ammonia (*52*), which were found elevated in feces of children with ASD (*48*). Genes identified as part of the pyruvate biosynthesis pathway were either found in both cohorts or only in predicted genomes associated with biomarkers from the NT cohort (Figure 2) (SI 9 panel B).

**Propionate pathway:** The predicted pathways for the synthesis of propionate, another SCFA, appears depleted in the ASD cohort (Figure C). This pathway has the potential to generate isopropanol, which has recently been found at a significantly greater concentration in the feces of children with autism (*43*).

**Sulfur Pathway:** The association between the predicted sulfur pathway and the NT cohort, though somewhat surprising, parallels our finding of *Acinetobacter* and *Desulfovibrio* enrichment in the NT cohort. The reactions detailed in SI 9 panel D suggest an imbalanced within the sulfur cycle, which has already been hypothesized as possible route modulating the gut-brain interaction in autism (*53*).

**Aminoacyl-tRNA Biosynthesis Pathway:** As expected, the vast majority of the aminoacyl-tRNA biosynthesis pathway is predicted to be present in all identified ESVs. Some enzymes from this pathway, specifically L-glutamine amido-ligase directly affect the availability of neurotransmitter precursors (SI 9 panel F) (*54*).

**Phosphotransferase system:** We also observed differential abundance of predicted carbohydrate uptake pathways within the ESVs associated with each cohort: the ESVs associated with ASD seemed to show a much greater variety of carbohydrate transporters (SI 9 panel G).

**Microbial metabolism in diverse environments:** This high level category comprises many different pathways, which were not individually found enriched in either cohort. It is however interesting to note that this KEGG category is associated with several metabolites already known in the literature as enriched or depleted in subjects with ASD (Figure 2). Among them were p-cresol (*43*), and ammonia (*49*) that have been found in greater abundance in the feces of children with autism; SCFAs (propanoate and acetate), which have mixed reports associating them with either children with ASD or controls (*9*, *49*, *55*); and neurotransmitters such as L-glutamate and GABA, which tend to be respectively greater and lower in feces of children with ASD, respectively (*43*). Glutamine, found in greater levels in the plasma of ASD participants (*43*), also belongs to this KEGG pathway, as do several metabolites such as nicotinate and aspartate, another neurotransmitter (*43*, *49*, *56*).

Et al

### Power Calculation and Sample Size

Likelihood-ratio-test statistics for a Dirichlet-Multinomial parameter test comparison showed that to reach an acceptable power (>0.9), we needed to include a minimum of 45 child-subjects per cohort. We were successful at screening, recruiting, sequencing, and analyzing 60 ASD and 57 NT child-subjects (SI 9). While this analysis does not calculate the power for each of the tests we performed, the Dirichlet-Multinomial distribution does allow power calculations for experimental design and population parameter estimations using a fully parametric approach. And though we cannot relate the verification of sufficient sample size to the non-parametric permutation test, we can conclude that our sample size is sufficient for reproducibility in the results from the zero-inflated Gaussian and DESeq2 models.

### Limitations

While these results show promising microbial differences between autism and typically developing children, potential limitations included reliance on self-reported information, limited identification of species or strain level variants, limited single time-point sampling, and lack of consideration of host genetic variation.

While we safeguarded against self-report bias through two validated machine-learning algorithms that adapt well to mobile testing, there remains bias in self-report of diagnosis may remain. In particular because we only required MARA for the child with ASD, we could not confirm the typical development of their siblings. In addition, the compliance with the optional request for video was slightly under <50% of the cohort studied. While it was encouraging to see perfect alignment between the MARA and the video classifier outcomes, bolstering confidence in the confirmation of self-report, it would be better to require this dual check for all participants in future work.

Although widely used (*19*), self-reported GI symptoms can also suffer some discrepancies when compared to a pediatric gastroenterologist reported data (*57*). Furthermore, while we observed physiological distinctions between the microbiomes of the cohorts on the level of exact sequence variants, it was often not possible to assign a taxonomic annotation or full genome to these sequences because of incomplete coverage in public databases. As the predicted pathways discussed were highly dependent on availabilities of full genome information, further metagenome and multi-Omics analyses in this space will be needed to confirm the metabolic hypotheses presented.

Finally, this study only collects one microbiome sample from each participant child and does not consider the influence of genetic variation between subjects or cohorts. A prospective and longitudinal study, described in Future Work, will ameliorate these limitations and significantly contribute to our understanding of the gut-brain interactions by accounting for the host genotype, gut microbiome, phenome, and metabolome of more than 200 age-matched sibling pairs with and without ASD.

### Future Work

This study has provided potential microbial biomarkers, both taxonomic and functional that associate the stool-associated microbiome with the ASD phenome. These findings may be due in part to the fact that we were able build a larger sample than previous studies on the microbiome and ASD, however, it will be necessary to continue sampling this modality in a larger and even more diverse cohort. Our study to confirms that crowdsourcing is a viable and cost effective way to do so. Thus our future work will take a similar angle but on an expanded population and will additionally move from single to multi-time-point sampling of the subjects to control for unrelated environmental influences on the gut microbiome. In addition, future work should combine the microbiome and phenotype, with the genome to enable more precise stratification of the relationship between microbiota and the Autism Spectrum and improve our understanding of the host-microbiome interaction as well as facilitate the discovery of more clinically useful autism biomarkers. This study phase has paved the way towards creating a more robust study where we will incorporate the three aforementioned modalities. We will also aim to validate whether specific microbial taxa and metabolisms are causally associated with ASD through improved characterization of the microbiome (e.g. metagenomics, metabolomics, etc.) in human longitudinal studies, animal studies to demonstrate causation, and human interventional studies that target the microbiome

### Conclusion

The aim of this study was to explore the association between the composition of the gut microbiome and the ASD phenotype in order to predict mechanism of association and to identify taxonomic and functional biomarkers and targets for future therapeutic research. Using a novel crowd-sourcing approach, we recruited 71 ASD/NT young sibling pairs, thereby limiting the confounding factors of age, lifestyle, diet, and genetics. This improves our confidence in the observed differences in gut bacterial taxa associated with the ASD phenotype. We observed systematic differences in the abundance of specific microbes between the ASD and NT cohorts, including a depletion of five ESVs from the Lachnospiraceae family and two ESVs associated with the genera *Desulfovibrio* and *Bifidobacterium* in the ASD cohort, and a difference in membership of clostridial organisms between the cohorts. Taxonomic assignments for short 16S rRNA fragments cannot be used to predict the full genetic functional potential of a microbiome, but using conservative methods we predicted the general functional potential of these microbiota and observed possible differences in the pathways associated with butyrate synthesis, potential harmful by-product and asssociated neurotransmitter production, that warrants further examination. The observed predicted differences in stool-associated microbial metabolic potential between ASD and NT siblings are worthy of future investigation into causality, and could represent opportunities for therapeutic intervention.

## MATERIALS AND METHODS

### Crowdsourcing Recruitment and Data Collection

Data were collected from March 2015 to September 2017 under an approved Stanford University Institutional Review Board protocol (eProtocol 30205). To target and inform the autism community of the study, we crowdsourced study subject recruitment via popular social media networking platforms including Twitter, Facebook, autism-focused Yahoo Groups, and a press released article from National Public Radio. In addition, we engaged with non-profit and for-profit companies who informed their community of the study via their social media platforms and email lists.

Parents of eligible participants completed the online component of study procedures via a secure, HIPAA-compliant web-based platform (https://microbiome.stanford.edu) where they provided electronic consent, responded to behavioral, demographic, and dietary surveys on behalf of their children, and uploaded a home video of the child with autism. Metadata were collected using RedCap (*58*, *59*).

### Sampling Kits

After they completed the online surveys, research staff mailed sampling kits to families for at-home stool collection. Each sampling kit included two sets of collections tubes and swabs to collect stool samples, for both the child with ASD and his or her neurotypically developing sibling, instructions on how to collect the samples, and a detailed, 53-question dietary questionnaire for each child (see Supporting Information SI 11). Participants returned the samples to the research staff via prepaid and pre-labelled packaging (*60*).

### ASD Diagnosis Confirmation

To confirm the parent-provided ASD diagnosis of a child-subject, we applied two machine learning classifiers, one based on a parent-directed questionnaire (*29*, *32*) and one based on a home video of the child with ASD (*28*, *29*, *33*). The parent-directed questionnaire is described below as “Mobile Autism Risk Assessment or MARA” and the video-based classifier is referred to as “video classifier. Due to both privacy and technical barriers to video upload, we made the video upload optional but required that all subjects complete the MARA as a strict inclusion criterion. We confirmed the self-reported diagnosis of the child using MARA and, when available, both the MARA and the video classifier. When both were available, concordance in outcome from both classifiers with the self-reported diagnosis was required for a sample to be included in our study.

### Mobile Autism Risk Assessment

Participants electronically completed the clinically validated Mobile Autism Risk Assessment (MARA) (*32*) (*29*). This system uses a set of 7 behavioral features developed through machine learning for rapid screening for autism. The 7-feature set is measured through parent-report in a questionnaire on a mobile device. Each feature is scored by the parent on a scale from 0 to 4, 0 being most impaired and 4 being least impaired. The features focus on the child’s language ability, make-believe play, social activity, restricted and repetitive behaviors, general signs of developmental delays by or before age 3, and eye contact. The responses generate a score that classifies the child as either “ASD” (positive score) or “no ASD” (negative score).

### Video Analysis

In addition, we requested (as optional) a home video of their child with ASD via our secure study website. For a video to be eligible for analysis, we asked that it include social interaction, use or play with objects in the video, be at least two minutes long, and clearly show the child’s face and hands. The specific details of scoring and the validation of the classifier are described in previous publications (*28*, *29*, *33*). For the purposes of this study we had at least 3 video raters who were blind to diagnosis independently tag the specific behavioral features that our video classifier requires to produce a risk score. We took the majority consensus diagnosis as the outcome for comparison with the caregiver’s self-reported diagnosis.

To safeguard against ascertainment bias due to increasing familiarity, we required our video analysts to score an unlabeled mixture of ASD participant videos and similar home videos of neurotypical children ages 2-7 years mined from YouTube’s publicly available video repository. The responses to each question were scored on a scale from 0 to 4, generating a classification of the child as either “ASD” or “no ASD”. Similar to the MARA, the outcome is a probability score that indicates both class as well as severity of phenotype.

### Lifestyle and Dietary Practices

Participants electronically completed dietary and lifestyle questionnaires on behalf of their children, using a 5-point frequency-based Likert scale and categorical answers (Supplementary Information SI 11). Questions covered dietary habits (e.g., How many servings of vegetables does your child eat in a typical week?), lifestyle habits (e.g., How many times a week does your child exercise?), and other pertinent information (e.g., Was your child born by C-section?).

We investigated systematic differences in the dietary habits of lifestyles of children with ASD as compared to NT children. Categorical data items were assigned either 1 or 0, and Likert scale items were assigned a value from 1 to 5 (1 = “Never,” 2 = “Occasionally,” 3 = “Sometimes,” 4 = “Often,” and 5 = “Always”). Differences of qualities with ordinal values were investigated using a linear-by-linear association test, and qualities with categorical values were tested using a chi-squared test. To verify that family relation was a practical criterion to ensure similarity between case and control lifestyle and dietary habits, we performed a permutation test (999 permutations) on the Euclidean distances between participants’ numerical responses. Data were standardized to mean as 0 and variance as 1 to account for the differences in scale between categorical and Likert scale numerical values.

### DNA Extraction, Amplification and Sequencing

Microbiome samples were processed according to the procedures outlined by the American Gut Project Protocol Apprill:2015gb, (*61*-*63*). DNA was extracted using the 96-well Powersoil DNA Isolation Kit (MO BIO, Carlsbad, CA). We utilized the manufacturer’s protocol with the following modification: after the addition of the sample and solution C1, we partially submerged the sealed extraction plates in a water bath for 10 minutes at 65°C. We amplified the extracted DNA using the 5PRIME MasterMix (5 PRIME, Inc, Gaithersburg, MD) and the 515F/806R primers for a final concentration of 0.2μM per primer. Thermocycler settings for generating amplicons were 3 minutes at 94°C, then 35 cycles at 94°C for 45 seconds, 50°C for 1 minute, and 72°C for 1.5 minutes, with a final extension for 10 minutes at 72°C. After PCR, we quantified the DNA concentration of each sample using the Quant-iT PicoGreen dsDNA Assay kit and then pooled to 70 ng DNA per sample. We generated clean pools using the QIAquick PCR Purification Kit (QIAGEN, Hilden, Germany). The clean pools were then submitted to the Environmental Sample Preparation and Sequencing Facility at Argonne National Laboratory to be sequenced on an Illumina MiSeq using V4 chemistry.

### Sequence Filtering, Chimera Removal, Taxonomic Assignment and Phylogenic Tree

Raw sequences were processed using the workflow available in the software package DADA2 (*41*), which models and corrects amplicon errors. Reads were trimmed to include base pairs 10 through 140 and truncated at the first instance of a Phred quality score less than 20. Reads with more than two expected errors were filtered out. Reads were then de-replicated and de-noised. Forward and reverse reads were merged and chimeras were removed. Taxonomy was assigned to each Exact Sequence Variant (ESV) generated by this pipeline by running the Ribosomal Database Project’s (RDP) naive Bayesian classifier (*64*), implemented in DADA2, against the GreenGenes dataset maintained in DADA2 package (*65*).

The phylogenetic tree was rooted using an archea sequence from *Halorhabdus rudnickae* as an outgroup (see available github code): all the sequences were aligned using the phangorn package and a Neighbor-Joining Tree was built (*66*) using ape. The tree was bootstrapped 100 times with phangorn (*67*).

### Statistical Analyses

We performed statistical analyses with R version 3.4.2 (2017-09-28) using R Studio Integrated development environment for R v1.0.136 (open source software, Boston, MA). We used the following packages in R: DESeq2_1.18.0, SummarizedExperiment_1.8.0, DelayedArray_0.4.1, matrixStats_0.52.2, GenomicRanges_1.30.0, GenomeInfoDb_1.14.0, IRanges_2.12.0, S4Vectors_0.16.0, BiocInstaller_1.28.0, gtable_0.2.0, cowplot_0.8.0, lattice_0.20-35, gridExtra_2.3, scales_0.5.0, metagenomeSeq_1.20.0, RColorBrewer_1.1-2, glmnet_2.0-13, foreach_1.4.3, Matrix_1.2-11, limma_3.34.0, Biobase_2.38.0, BiocGenerics_0.24.0, gage_2.28.0, readr_1.1.1, igraph_1.1.2, ggplot2_2.2.1, reshape2_1.4.2, structSSI_1.1.1, dplyr_0.7.4, ape_5.0, phyloseq_1.22.3. All code used for this work is publicly available: https://github.com/walllab/ASD_microbiome16s_public/. The raw fastq files can be found at (*13*, *43*). For a workflow diagram, see SI 12.

### Analysis of Alpha-Diversity Differences

We calculated alpha-diversity for each sample using Shannon-Weiner diversity, a traditional metric that takes into account richness and evenness of taxonomic species, and Phylogenetic Diversity, a metric that measures the total length of phylogenetic branches necessary to span the set of taxa in a sample (*39*). We then used a Wilcoxon rank sum test (*68*) to quantify the significance of differences observed between the two cohorts. Next, we performed 1000 bootstrap simulations to calculate the variance of diversity metrics observed in each cohort and again used a rank sum test to quantify significance of the difference in variances.

### Identification of Dietary and Lifestyle Habits Influencing the Microbial Community

To determine whether or not the parent-reported dietary and lifestyle questionnaires contained influential data and insight into the microbial communities observed in our samples, we used a PERMANOVA test (ADONIS function in vegan package) on Bray-Curtis distances (*13*). We also performed a test to measure the homogeneity of the dispersion (PERMDISP2 procedure) of each cluster in order to be more confident that the cluster-specific centroids were robustly different rather than due to disparities in cohort dispersions (*19*).

### Permutation Test on Sibling Pair Differentials

In addition to community level trends, we investigated differential abundances of specific taxa. We ran a permutation test on mean taxa abundance differences between sibling pairs to determine if any taxa were systematically enriched or depleted in our ASD samples when compared to our NT controls. We selected DeSeq2 as a normalization method to minimize batch effect and noise from differences in sampling depth while maintaining the expected dataset properties (*69*). We expected that the microbiome compositions of age-matched sibling pairs would be closer to each other than to any other samples in the cohort, due to the environmental and genetic similarities shared by young siblings living within the same household. Additionally, we resampled and resequenced samples from eight individuals, with at least six months in between samplings, to examine the similarities of the results and confirm that the quality of the samples was maintained over time. Just as in the case of siblings, we expected samples from the same individual to contain microbiome compositions very similar to each other. We investigated the suitability of three commonly used normalization techniques: CSS normalization, DeSeq2 normalization, and log transformation, and found that DeSeq2 best matched above stated expectations (SI 13). This method performs variance stabilization to normalize counts with respect to library size and heteroskedasticity (*70*).

We first excluded all samples that were singletons resulting from quality control or from third siblings (sibling furthest in age was removed). We were left with 55 sibling pairs, each sequenced on the same day, on the same instrument and to similar depths. Using DeSeq2 normalized taxa abundances, counts in the NT siblings were subtracted from those in the ASD siblings and averaged across all samples for each taxon. For each taxon, we simulated a null distribution of the average sibling differences by repeating the above procedure with permutated sibling pairs and phenotype assignments 10000 times. Lastly, we calculated a p-value as the number of times the null hypotheses produced a value more extreme than the actual value observed (see SI H).

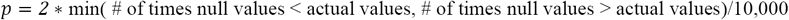

Null distributions remained stable well before 10000 permutations. We assessed stability by comparing the value of the ks test statistic to the null distribution shapes at increments of 500 simulations. At 10,000 simulations, the maximal ks test statistic over all taxa (when increasing from 9000 to 10000 simulations) was 4.8 * 10^-3.

### Models to Maximize the Likelihood of Detecting Low Abundance Species

Given the sparsity of 16S sequencing due to sequencing depth limitations, we used differential ribosomal analysis based on the negative binomial distribution and zero inflated Gaussian analysis, to estimate log-fold changes of taxa abundances between our ASD and NT groups (*71*).

### Differential Ribosomal Analysis Based on the Negative Binomial Distribution

We performed differential analysis of taxa counts between groups by modeling taxa abundances under a negative binomial model using the DESeq2 framework. This method performs variance stabilization on taxa counts and then fits a generalized linear model with a log link on normalized count data. Coefficients representing the log fold changes of taxa between groups are then extracted and shrunk toward zero using an empirical Bayes model that effects taxa with low counts more severely. The method then calculates each shrunken log fold change’s standard error from the curvature of its posterior and performs a Wald test (*72*) to determine whether the log fold change of any one taxa is significantly different from zero. The p-values associated with each taxon are corrected for multiple hypotheses using false discovery rate (FDR) (*73*).

### Zero Inflated Gaussian Analysis

To account for sparsity due to under-sampling, we used the method developed by Paulson *et al.*: a mixture model that uses a zero-inflated gaussian distribution to account for varying depths of coverage (*35*). To model data appropriately under a zero-inflated Gaussian model, it was necessary to normalize data in a way that does not change the distribution of the variance of taxa across samples. We used cumulative sum scaling (CSS) (*35*) to account for under-sampling and increase the sensitivity and specificity of identifiable taxa. CSS is a technique that mitigates bias coming from features that are preferentially amplified in a sample-specific manner. CSS divides the feature counts for each sample by the sum of feature counts with values less than that of the median (or chosen percentile), rather than by the total counts in that sample (as in total-sum normalization). Counts are then multiplied by a normalization constant that is the same across samples to ensure normalized counts have interpretable units. The method models each taxon count as a mixture model of a point mass at zero and a normal distribution parameterized by the observed taxa distribution.

### Functional Profile Prediction using Piphillan

We inferred metabolic activity using Piphillan (*47*), a bioinformatics software package designed to predict metagenome functional content from marker gene (16S) surveys. Piphillan aligns 16S sequences to sequences in the GreenGenes database (*65*), and assigns functional profiles based on 97% match of 16S sequences. 16S sequences that do not match a database entry are assigned the functional profile of their nearest neighbor. We chose this software over other alternatives, such as PiCRUST (*74*) and Tax4Fun (*75*), because of its high performance on clinical samples and its usage of the most current GreenGenes database. Piphillan was run on DESeq2 normalized data to produce estimates of KEGG ortholog abundances (*76*-*78*). We then performed a modified gene enrichment analysis: a set was considered a metabolic pathway as defined by the KEGG Brite database and an element was considered a KEGG ortholog (*79*).

### Gene Set Enrichment Analysis Using KEGG Orthologs

Using KO prediction provided by Piphillan, we distilled the data to define functional pathways as described by the KEGG database <http://www.kegg.jp/kegg/pathway.html> (*78*). As some KOs contribute to multiple pathways, we first divided the abundance of each KO in a sample by the number of pathways the KO participates in, so that a single functional unit’s activity or output may contribute to only one pathway at a time (as a rule of thumb). Then we used the Gage implementation of Gene Set Enrichment Analysis (GSEA) (*80*) to perform a modified gene enrichment analysis: a set was considered a metabolic pathway as defined by the KEGG Brite database and an element was considered a KEGG ortholog(*79*).

### Power Calculation

We modeled microbial abundances as a Dirichlet-Multinomial, a model which has been proven to successfully reflect the abundances seen in naturally occurring microbial communities (*81*). Under this model, we estimated Method-of-Moments (MoM) parameters for each taxon and determined the stability of those estimates by comparing likelihood-ratio-test statistics over 1000 Monte-Carlo simulations (*82*). From this simulation, we can determine the number of samples necessary to reach a given level of power when estimating parameter values. At a rejection threshold value of 0.05, n = 70 ASD child-subjects and n = 70 NT subjects were required to obtain power above 0.99, and n = 45 ASD child-subjects and n = 45 NT child-subjects was sufficient to provide a power greater than 0.95. Though we do not explicitly use the Dirichlet-Multinomial model in further analyses, a high power in this context implies a high power in more complex down-stream analyses that are not able to be simulated.

## Acknowledgements

The work was supported in part by funds to DPW from NIH (<u>1R01EB025025-01</u> & <u>1R21HD091500-01</u>), The Hartwell Foundation, Bill and Melinda Gates Foundation, Coulter Foundation, Lucile Packard Foundation, and program grants from Stanford’s Precision Health and Integrated Diagnostics *Center* (*PHIND*), Beckman Center, Bio-X Center, Predictives and Diagnostics Accelerator (SPADA) Spectrum, and Child Health Research Institute. We also acknowledge generous support from David Orr, Imma Calvo, Bobby Dekesyer and Peter Sullivan.

